# Adolescent tuning of association cortex in human structural brain networks

**DOI:** 10.1101/126920

**Authors:** František Váša, Jakob Seidlitz, Rafael Romero-Garcia, Kirstie J. Whitaker, Gideon Rosenthal, Petra E. Vértes, Maxwell Shinn, Aaron Alexander-Bloch, Peter Fonagy, Raymond J. Dolan, Peter B. Jones, Ian M. Goodyer, the NSPN consortium, Olaf Sporns, Edward T. Bullmore

**Author notes:** List of NSPN consortium members in Supplementary Information. Correspondence to: František Váša, Brain Mapping Unit, Department of Psychiatry, Sir William Hardy Building, Downing Street, Cambridge CB2 3EB.

## Abstract

Motivated by prior data on local cortical shrinkage and intracortical myelination, we predicted age-related changes in topological organisation of cortical structural networks during adolescence. We estimated structural correlation from magnetic resonance imaging measures of cortical thickness at 308 regions in a sample of N=297 healthy participants, aged 14-24 years. We used a novel sliding-window analysis to measure age-related changes in network attributes globally, locally and in the context of several community partitions of the network. We found that the strength of structural correlation generally decreased as a function of age. Association cortical regions demonstrated a sharp decrease in nodal degree (hubness) from 14 years, reaching a minimum at approximately 19 years, and then levelling off or even slightly increasing until 24 years. Greater and more prolonged age-related changes in degree of cortical regions within the brain network were associated with faster rates of adolescent cortical myelination and shrinkage. The brain regions that demonstrated the greatest age-related changes were concentrated within prefrontal modules. We conclude that human adolescence is associated with biologically plausible changes in structural imaging markers of brain network organization, consistent with the concept of tuning or consolidating anatomical connectivity between frontal cortex and the rest of the connectome.

---

Human adolescence is known to be a major phase of cortical development. In particular, cerebral cortex becomes thinner (Wierenga et al., 2014) and more densely myelinated (Miller et al., 2012) in the transition from puberty to young adulthood. Adolescent decreases in cortical thickness (thinning) are variable between different areas of cortex (Raznahan et al., 2011): for example, thinning is greater in association cortical areas than primary sensory areas (Whitaker, Vértes et al., 2016).

Motivated by these and other results, we predicted that human adolescence should be associated with changes in the architecture of structural brain networks. There are currently only two experimental techniques, both based on magnetic resonance imaging (MRI), that are capable of providing data to test this prediction: diffusion tensor imaging followed by tractography; or structural MRI followed by structural covariance or correlation analysis. Here we focused on the latter, measuring the thickness of a set of predefined cortical regions in each individual MRI dataset and then estimating the correlation of thickness between each possible pair of regions across participants. Similar methods have been widely used and validated (Lerch et al., 2006) in a range of prior studies (Alexander-Bloch et al., 2013; Evans, 2013).

In particular, structural correlation (covariance) measures have been used as a basis for graph theoretical modelling of the human connectome (Bullmore & Sporns, 2009; Fornito et al., 2016). Considerable evidence has accumulated in support of the general view that human brain structural correlation networks have a complex topological organization, characterised by non-random features such as the existence of highly connected (high degree) hub nodes and a modular community structure (Alexander-Bloch et al., 2013; Evans, 2013). Topological metrics on structural correlation networks have demonstrated changes associated with disease, development and ageing (Alexander-Bloch et al., 2013; Evans, 2013). However, only two studies have investigated adolescent changes in structural correlation networks. Zielinski et al. (2010) demonstrated that the anatomical extent of structural correlation networks, assessed using seed-based correlation of voxel-wise grey matter intensity, changes in adolescence in a spatially patterned manner. Specifically, primary visual and sensori-motor networks, as well as the default mode network, expanded in early childhood before being "pruned" in adolescence, while higher-order cognitive networks showed a gradual monotonic gain in spatial extent. Subsequently, Khundrakpam et al. (2013) applied graph-theoretical analyses to a subset of the same data, reporting childhood increases in topological integration (global efficiency) and decreases in topological segregation (local efficiency and modularity), as well as increases in regional integration in paralimbic and association regions. While these studies constitute interesting initial investigations, their ability to precisely describe developmental changes is limited by their segregation of participants into four discrete age-defined strata, resulting in relatively coarse-grained resolution of brain maturational trajectories.

Here, we aimed to obtain a more precise description of adolescent maturational trajectories of structural network architecture, which were hypothesised to vary as a smooth and potentially non-linear function of age. We used a sliding-window analysis to estimate structural correlations and structural network properties for each of an overlapping series of 9 age-defined windows or strata of the sample (N≈60 participants per window). We identified the cortical regions (nodes) and connections (edges) which showed the most significant age-related changes in structural correlation. We tested the related hypotheses that parameters of adolescent change in structural correlation would be greater and occur later in regions of association cortex, which show faster rates of local cortical shrinkage and myelination. In addition, we explored whether greater and later changes in structural correlation during adolescence would be concentrated within or between specific communities of regions. Specifically we mapped adolescent changes in structural correlation to three brain community structures: the topological modular partition of the age-invariant structural correlation network; an atlas of cytoarchitectonic classes (von Economo & Koskinas, 1925); and functional intrinsic connectivity or resting state networks (Yeo, Krienen et al., 2011).

## Materials and Methods

### Participants

A demographically balanced cohort of 297 healthy participants (149 females) aged 14-24 years was included in this study, with approximately 60 participants in each of 5 age-defined strata: 14-15 years inclusive, 16-17 years, 18-19 years, 20-21 years, and 22-24 years. Participants were excluded if they were currently being treated for a psychiatric disorder or for drug or alcohol dependence; had a current or past history of neurological disorders or trauma; or had a learning disability. Participants provided informed written consent for each aspect of the study, and parental consent was obtained for those aged 14–15 years. The study was ethically approved by the National Research Ethics Service and was conducted in accordance with NHS research governance standards.

### MRI acquisition and processing

Structural scans were acquired at three sites using multi-parametric mapping (MPM) implemented on three identical 3T MRI scanners (Siemens Magnetom TIM Trio). Inter-site reliability of the sequence was evaluated within a pilot study of five healthy participants each scanned at each site (Weiskopf et al., 2013). The MPM sequence includes maps of R_1_ (1/T_1_) and magnetization transfer (MT), indicative of myelination. For details of MRI acquisition parameters, see **Supplementary Information (SI)**.

Processing of individual scans using FreeSurfer v5.3.0 included skull-stripping, segmentation of cortical grey and white matter and reconstruction of the cortical surface and grey-white matter boundary (Fischl et al., 1999). All scans were stringently quality controlled by rerunning the reconstruction algorithm after the addition of control points and white matter edits (details in SI). The cerebral cortex of each participant was parcellated into 308 regions of interest, based on a sub-division of the Desikan-Kiliany anatomical atlas (Desikan et al., 2006) into parcels of approximately equal surface area (~5cm^2^) (Romero-Garcia et al., 2012).

Regional changes in cortical thickness (CT) and myelination (MT) were characterized using the rate of change over adolescence, evaluated as the slope of a linear model fitted to the crosssectional values. Following Whitaker, Vértes et al. (2016), myelination analyses were conducted at 10 fractional depths between the pial surface and the grey/white matter boundary, as well as two absolute depths into white matter. Main analyses focused on MT estimates at 70% fractional cortical depth from the pial surface. For details and results across cortical depths, see the **Supplementary Information**.

While both cortical thickness and myelination maps were averaged within parcels, for comparison between maturation of structural correlation networks and morphology, only the cortical thickness values were used to construct structural correlation networks.

### Age-invariant structural network

An age-invariant structural correlation network was constructed using Pearson correlations in cortical thickness between pairs of regions across all 297 participants, to serve as a reference for developmental changes within the age-resolved structural networks (described below; Fig. 1A). We used raw cortical thickness values, uncorrected for age, gender or intracranial volume. However, correcting for these covariates had no effect on the results. For background reading on graph theoretical methods and connectomics see Bullmore & Sporns (2009) and Fornito et al. (2016).

**Figure 1:**
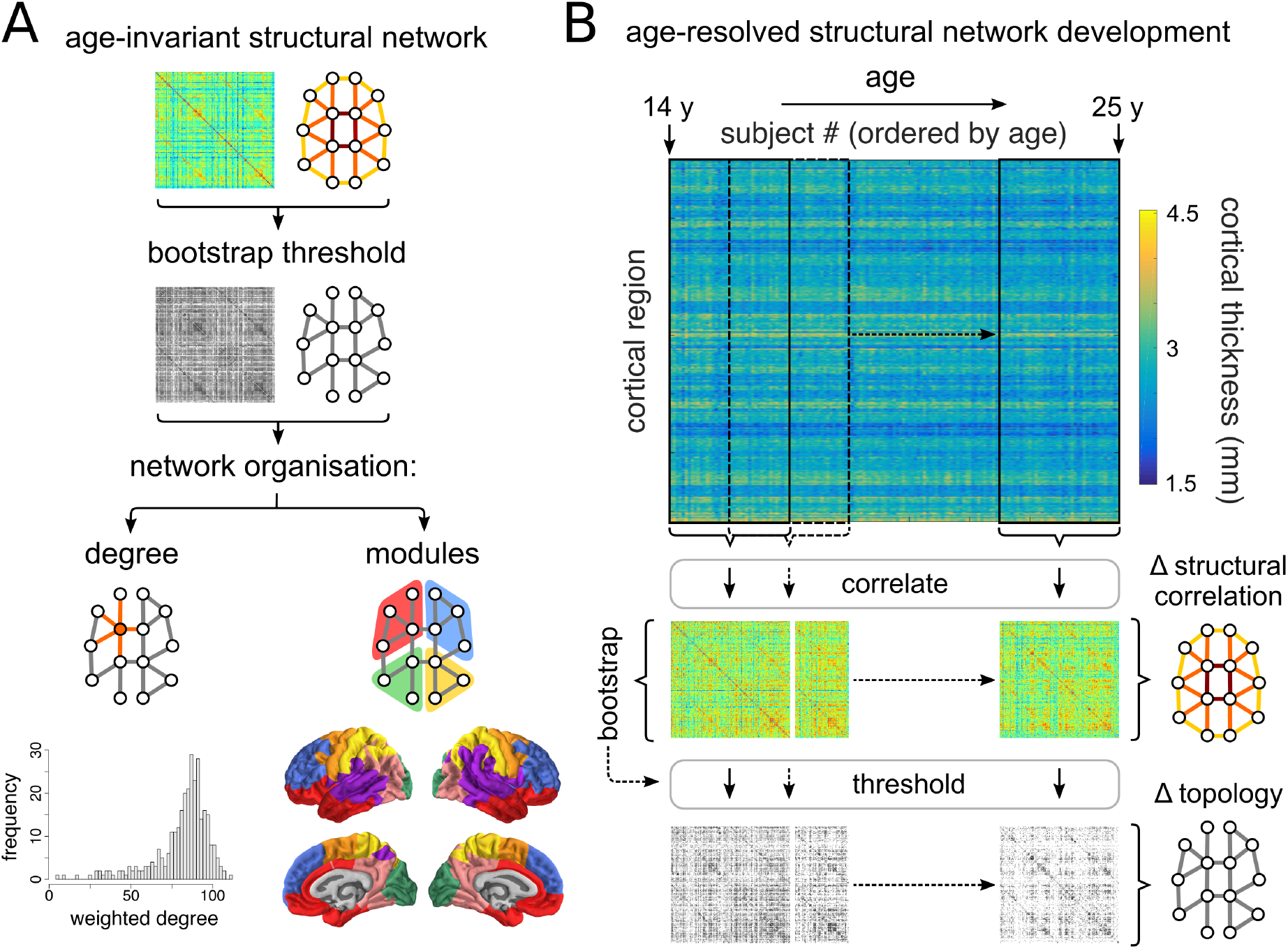
Construction of age-invariant and age-resolved structural correlation networks. A) An age-invariant structural correlation network was constructed by cross-correlating regional cortical thickness across all participants. This network was probabilistically thresholded using a bootstrap-based method. Network organisation was evaluated using several measures, including the degree (both binary and weighted; respectively the number and sum of weights of retained edges connected to a node) and modular architecture. For details regarding module generation, see supplementary information (**SI Fig. S1**). B) Age-resolved structural correlation networks were constructed using a sliding-window method. Participants were ordered by age, and structural networks were constructed by estimating correlations between regional cortical thickness values across participants within overlapping windows iteratively slid across the age range. Correlations were probabilistically thresholded using bootstrap, before developmental trajectories were fitted to summary window-derived measures as a function of the median age of participants within each window.

The age-invariant structural network was thresholded using a bootstrap approach, whereby 1000 sets of participants were resampled with replacement and used to construct surrogate structural networks. For each pair of regions, we examined whether there is evidence of a non-zero correlation across bootstraps: edges that were consistently positive or negative across bootstraps (at a two-tailed, FDR-adjusted level of α = 0.01) were retained; the remaining edges were set to zero. Nodal topological organisation of the thresholded network was assessed using degree, defined as the number of retained correlations for each node, as well as the weighted degree, or summed weight of retained edges for each node.

Further, the age-invariant network was partitioned into communities of nodes showing higher structural correlations within than between communities (Sporns & Betzel, 2016). The community structure of the age-invariant network was decomposed using the Louvain multiresolution algorithm (Blondel et al., 2008) over the resolution parameter range 0.01 ≤ γ ≤ 4.00. As γ increases, the community structure is decomposed to a progressively larger number of modules. We used the concept of minimizing versatility to identify those resolution parameter values which reduce the uncertainty with which any node was affiliated consistently to the same module (Shinn et al., 2017). The final community partition was defined as a consensus across 1000 runs of the Louvain modularity algorithm (Lancichinetti & Fortunato, 2012) at the selected value of the resolution parameter γ. For details regarding module generation, see **SI Fig. S1**.

### Development of age-resolved structural networks

#### Sliding window network construction

Development of structural networks between 14 and 24 years was evaluated using a sliding window method. Regional cortical thickness values were cross-correlated within windows containing equal numbers of participants, and incrementally slid across the age-range by regular increments (Fig. 1B). The two parameters of the method, the "window width" and the "step size" (in units of number of participants) determine the number of windows, each of which generates a structural correlation network. Exploration of the sliding window parameter values suggests that results are qualitatively consistent across a range of parameter combinations. For the (in)dependence of results on sliding window parameters, and a discussion of the considerations involved in parameter selection, see the **Supplementary Information**.

Results presented below correspond to nine half-overlapping windows of 60 participants each, obtained by interpolating the five age strata of the NSPN study, within which participants were recruited. Gender was relatively balanced within the interpolated bins, with the most imbalanced ratio being 34:26 = 57%:43% (M:F). We investigated the effects of gender separately (see below).

Global maturation of structural networks was characterised using the mean of the correlation distribution. At the regional level, an analogous measure was used – nodal strength, the mean of the pattern of regional correlations (rows, or equally, columns of the correlation matrices).

#### Bootstrap thresholding of age-resolved structural networks

Estimating structural correlation networks from a small number of participants is an inherently noisy process; therefore, our principal analyses focused on networks probabilistically thresholded using bootstrap (Fig. 2B). The bootstrap thresholding procedure was identical to the one described above for age-invariant networks, but in this case was applied within windows. From the set of participants included in each window, an equal number of participants was sampled with replacement and the correlation structure was reestimated 1000 times. For each pair of regions, we examined whether there is evidence of a non-zero correlation across bootstraps: edges that were consistently positive across bootstraps (at a two-tailed, FDR-adjusted level of α = 0.01) were retained (there were no consistently negative edges); the remaining edges were set to zero.

**Figure 2:**
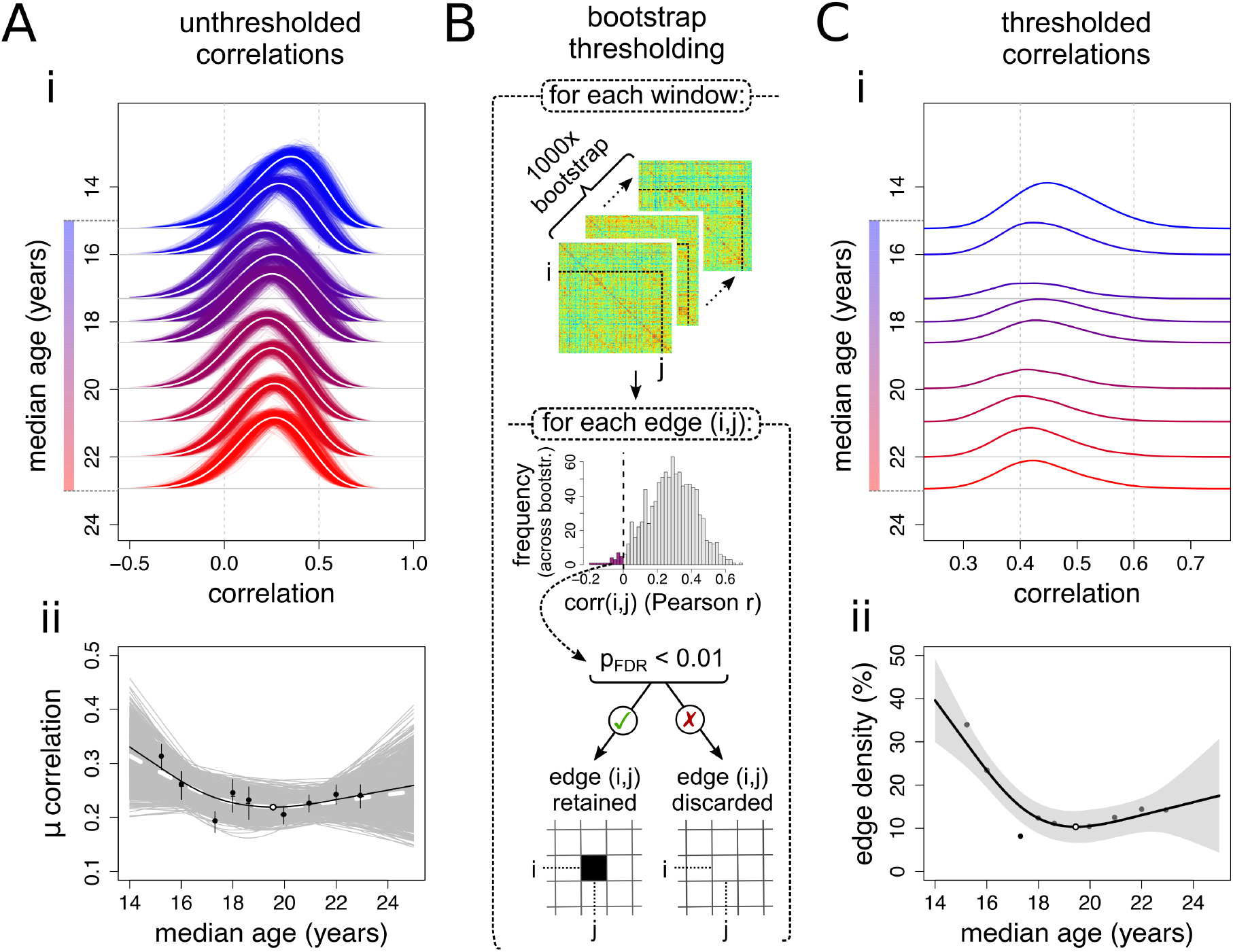
Global trajectories of age-resolved structural correlations and network connection density. A) Global trajectories of unthresholded structural correlations. (i) Development of the distribution of unthresholded correlations across age windows. Thin lines represent bootstrapped estimates, white lines represent the bootstrap mean. (ii) Changes in the average correlation. Black markers represent empirical data (error bars indicate the interquartile range across bootstraps), with corresponding regression line; the white marker indicates the trajectory minimum. Grey lines represent bootstrapped trajectories; the white dashed line represents the bootstrap mean. B) Each windowed matrix was thresholded using bootstrap. Within each window, 1000 sets of participants were resampled (with replacement) and used to construct correlation matrices. For each edge (correlation) within each window, the presence of a significant non-zero correlation (across bootstraps) was tested at the FDR-adjusted level of α_FDR_ = 0.01. Consistent correlations were retained, while inconsistent correlations were assigned a value of 0. C) Global trajectories within thresholded structural correlation networks. (i) Development of the distribution of correlations retained after probabilistic thresholding across age windows. (ii) The number of edges retained after probabilistic thresholding, or edge density. The shaded area represents the 95% confidence interval of the spline fit.

The global topological organisation of the thresholded graphs was assessed using the edge density, defined as the percentage of retained edges (relative to their possible total), as well as the distance spanned by retained edges, calculated as the average Euclidean distance between centroids of corresponding nodes. Nodal topological organisation was assessed using (analogous) measures of degree, defined as the number of edges connected to a node, and average Euclidean distance spanned by a node's retained edges. We have focused on simple graph-theoretical measures, such as edge density and node degree, for two reasons: first, our bootstrap-thresholded networks display variable edge density, which many "higher-order" graph-theoretical measures show a strong dependence on (van Wijk et al., 2010), and (2) even in correlation-based networks thresholded to fixed edge density, graph theoretical properties display a dependence on more elementary statistics such as properties of the correlation distribution (van den Heuvel et al., 2017).

### Fitting and characterisation of developmental trajectories

Developmental trajectories were fitted to both global and local measures as a function of the median age of participants in each window. In addition to linear models, we fitted locally adaptive smoothing splines. The nonparametric smoothing spline was chosen to model nonlinear trajectories over parametric alternatives as it was shown to be superior to quadratic fits in studies of brain development (Fjell et al., 2010). Still, the spline fits were constrained to be (approximately) at least as smooth as a quadratic fit (i.e.: effective degrees of freedom, *df* ≤ 3.5), based on the hypothesis that adolescent developmental trajectories over a 10-year age range should not display greater complexity. The specific smoothing spline used was a weighted sum of 6 cubic b-splines with knots placed at quantiles of the data and smoothing optimised using restricted maximum likelihood (REML) (Reiss et al., 2014). The relative quality of linear and spline fits, given their parsimony, was assessed using Akaike's information criterion (AIC). Classification using the Bayesian Information Criterion (BIC) yielded consistent results.

Regional changes were summarised using measures of maximum change in degree Δ*k*_max_, quantified as the difference between maximum and minimum degree, and the age at minimum degree age(*k*_min_). Further, we classified regional changes in degree as linear or nonlinear (using the AIC), and as increasing or decreasing (using the direction of maximum change). As an alternative measure of the magnitude of regional changes in structural correlation, we extracted linear rates of change of degree; the results were qualitatively consistent with the measure of maximum change, which is more suitable for non-linear trajectories (**Supplementary Information**).

### Relationship of structural network development to age-invariant network architecture

Given our previous finding, that highly correlated "hub nodes" of the age-invariant structural network (derived from all participants) are regions which thin and myelinate most over adolescence (Whitaker, Vértes et al., 2016), we were interested in studying the relationship of structural network development to age-invariant structural network architecture.

We evaluated Spearman's rank correlations between node degree in the age-invariant structural network, and parameters of change in node degree within the age-resolved structural network – including the amplitude of maximum change in degree Δ*k*_max_ as well as the age at minimum degree age(*k*_min_).

Finally, we studied changes in structural network organisation relative to three sets of node communities, including the partition of the age-invariant network into modules, the von Economo atlas of cytoarchitectonic classes (von Economo & Koskinas, 1925), and a set of functional intrinsic connectivity networks (Yeo, Krienen et al., 2011). For each community template and each age-window, we calculated the density of edges *D* within each community as well as between each pair of communities (within the same template), as the ratio of existing edges relative to the maximum number of possible edges in this within or between-community edge set. We then characterised changes in edge density within and between communities using measures analogous to the nodal trajectories – maximum change in edge density Δ*D*_max_ and age at minimum density age(*D*_min_). For details regarding the matching of the community templates to our 308-region parcellation, see the **Supplementary Information**.

### Spatial permutation test

In several analyses in the current study, measures were related to each other across regions. While numerous studies have reported significance based on the assumption that the number of samples is equal to the number of regions, this is technically inaccurate, as the number of regions is both arbitrary (due to the resolution of the chosen parcellation) and nonindependent (due to spatial auto-correlation amongst neighbouring parcels). To address this issue, spatial permutation tests have been implemented in past studies (Alexander-Bloch et al., 2013; Vandekar et al., 2015), which consist in comparing the empirical correlation amongst two spatial maps to a set of null correlations, generated by randomly rotating the spherical projection of one of the two spatial maps (as generated in FreeSurfer or Caret) before projecting it back on the brain surface. Importantly, the rotated projection preserved spatial contiguity of the empirical maps, as well as hemispheric symmetry. Such tests were previously implemented at the vertex level (Alexander-Bloch et al., 2013; Vandekar et al., 2015; here we implemented an analogous permutation test at the regional level. Thus, each analysis correlating values from two cortical maps is reported with both the p-value corresponding to the Spearman correlation (p_Spearman_), as well as a p-value derived from the spherical permutation (p_perm_), obtained by comparing the empirical Spearman's *ρ* to a null distribution of 10'000 Spearman correlations, between one empirical map and randomly rotated projections of the other map. For full details on the spherical permutation test, see **Supplementary Information**.

### Sensitivity analyses

To ascertain the robustness of obtained results to sliding window parameters and other methodological decisions and to rule out effects of potential artefactual causes, we conducted several ancillary studies.

We first investigated effects of sliding window parameters by systematically varying the window width and step size over ranges of {40,60,80} and {5,10,20} participants respectively.

Further, we examined potential effects of gender by repeating sliding window analyses separately for each gender (149 female, 148 male participants). This resulted in nine windows of ~30 participants each. Following estimation of global and nodal sliding window statistics separately for each gender within both unthresholded and bootstrap-thresholded networks (as described for all participants above), we fitted linear and spline models to the combined data, separately modelling effects of age, gender, and the age-by-gender interaction.

Finally, we studied the effect of several potential artefacts, including the presence of regions with low reliability of structural correlations as well as irregularities in the age distribution of participants.

For full results and discussion of these additional studies, see **Supplementary Information**.

## Results

### Age-invariant structural network

We first considered the structural correlation network constructed by thresholding the pairwise inter-regional correlations estimated from cortical thickness measurements on all (297) participants, age range 14-24 years (inclusive). Since this analysis combines data from all ages in the sample, we can refer to the result as an age-invariant structural correlation network (Fig. 1A).

The distribution of structural correlation had a positive mean value and was approximately symmetrical. The structural correlation matrix was thresholded probabilistically, using a bootstrap-based resampling procedure (**Methods**), to control the edge-wise false positive rate. Since this thresholding operation entailed approximately 47,000 hypothesis tests, we used the false discovery rate (FDR) algorithm to adjust for multiple comparisons. The resulting graph was densely connected (connection density ≈ 90%) and exhibited a modular community structure (Fig. 1A). The community partition consisted of seven modules, including three primary cortex modules: somatosensory (anterior parietal cortex), motor (posterior frontal cortex) and visual (occipital cortex), as well as an inferior-frontal/temporal module, a superior frontal module, a superior temporal/insular module and a parieto-occipital module. For details on this community structure and other modular partitions comprising different numbers of modules see SI Fig. S1 and SI Table S1.

### Age-resolved structural networks

To resolve age-related changes in structural networks, we used a "sliding window" analysis to estimate the structural correlation matrix separately for each of a series of subsets of the sample defined by overlapping age ranges or windows (Fig. 1B). The results of this analysis are naturally somewhat dependent on the sliding window parameters: the age-range spanned by each window and the incremental step between windows. Below we focus on results obtained with 9 windows of ~60 participants each, ranging from [14.1-16.0 years] to [22.0-25.0 years] with an incremental step of 30 participants (~1 year). We also explored a range of alternative sliding window parameters and demonstrated that our key results were robust to this methodological variation (**Supplementary Information**.)

Globally, over the whole brain, there was a non-linear trend of reducing structural correlation from the youngest age window to the oldest age window (Fig. 2A). Relatively strong positive correlations at age 14 (>0.31) decreased sharply over the next few windows, with minimum mean correlation (~0.22) occurring at 19.59 years (95% confidence interval (CI) [19.37, 19.76] years) and then slightly increasing again towards age 24 (AIC_spl_ < AIC_lin_, r^2^_adj_ = 0.52, p = 0.098; Fig. 2Aii). Both the mean inter-regional covariance, and the mean product of regional standard deviations (respectively the numerator and denominator of the Pearson correlation coefficient), showed similar non-linear processes of decline in younger windows followed by levelling off in older windows (SI Fig. S4).

A potential drawback of the sliding window analysis is that it inevitably involves estimating inter-regional correlations on a subset of the sample (N≈60 per window), with commensurately reduced precision of estimation and therefore noisier graphs. We used a probabilistic threshold to control the edge-wise FDR at 1%, thus ensuring that the age-resolved graphs only included edges that were unlikely to represent false positive noise (Fig. 2B).

Focusing on the most statistically robust subset of edges (which passed the FDR threshold for significance), we found similar but clearer evidence for age-related global changes in structural network organisation. The structural correlation distributions of the bootstrap-thresholded network became sparser over the course of adolescence (Fig. 2Ci). The edge density demonstrated a non-linear decrease (AIC_spl_ < AIC_lin_) from 33.9% to a minimum of 8.2% at 19.45 years (95% CI [19.32, 19.59] years; r^2^_adj_ = 0.81, p = 0.0069), which was similar in shape to the global trajectory of unthresholded correlation (Fig. 2Cii).

The global connection distance of the thresholded networks (the mean Euclidean distance subtended by bootstrap-thresholded edges) also demonstrated a non-linear trajectory (AIC_lin_ < AIC_spl_, r^2^_adj_ = 0.67, p = 0.049) characterised by a phase of relatively rapid decrease from 14 years to reach a minimum at 18.72 years (95% CI [18.68, 18.77] years), followed by a phase of more stable connection distance (SI Fig. S7A).

### Regional development of age-resolved structural networks

Regional maturation of structural correlation networks was assessed by estimating the trajectories of changes in node degree, which is the number of correlations retained at each node (following bootstrap thresholding). Although there was regional heterogeneity in the trajectories of node degree (Fig. 3A), all regions that demonstrated significant evidence of non-zero change (linear or spline fit p_FDR_ < 0.05; 82 regions) followed a nonlinear trajectory (AIC_spl_ < AIC_lin_), which for most regions (75/82) could be summarised by a younger phase (from 14 to 19 years approximately) of more-or-less rapid decrease in structural correlation followed by a levelling off or slight increase of structural correlation in an older phase (from 19 to 24 years approximately). This process could be summarised by two parameters: Δ*k*_max_, the difference between maximum and minimum degree; and age(*k*_min_), the age at which node degree reached its minimum value (Fig. 3B).

**Figure 3:**
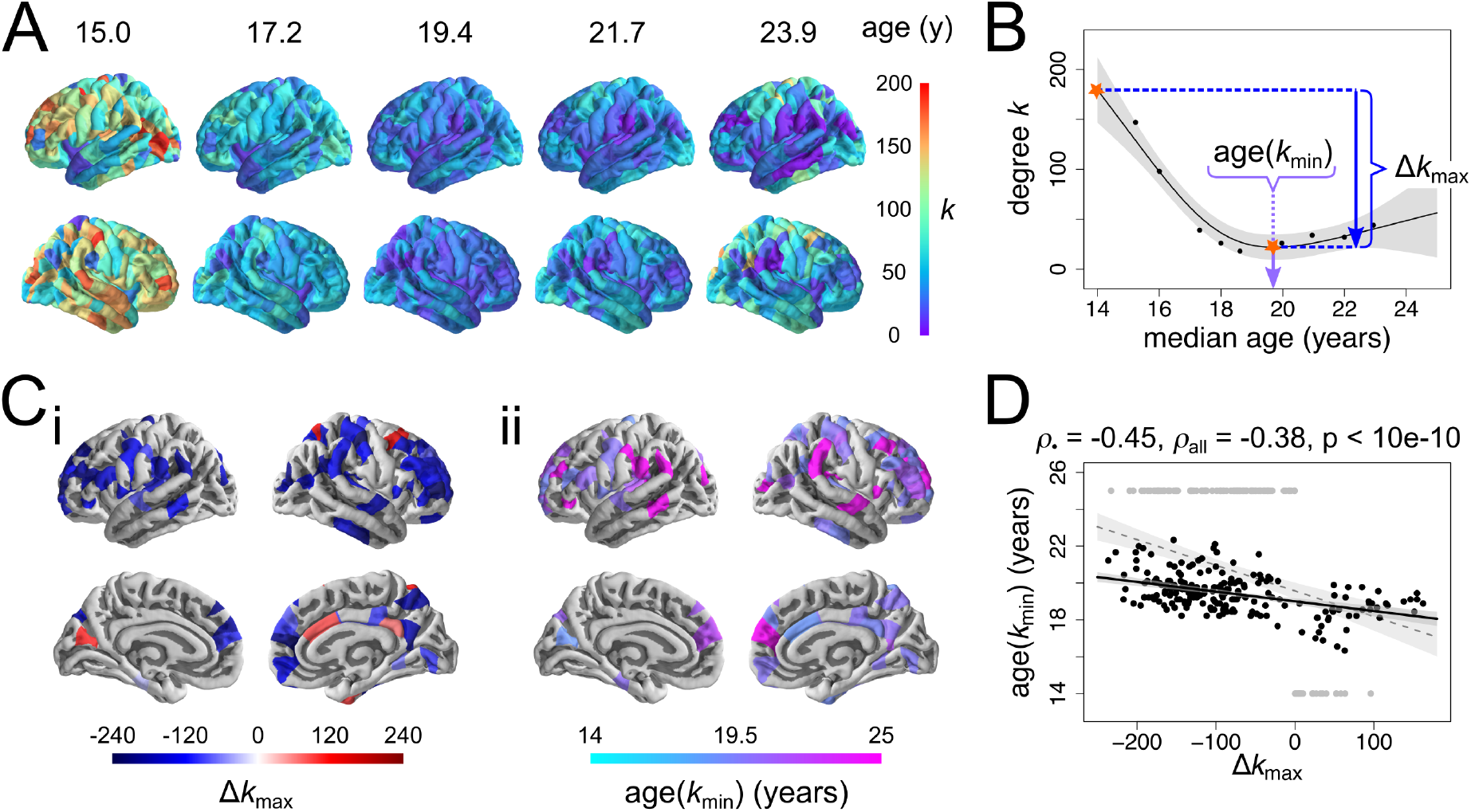
Regional development of structural correlation networks. A) Cortical maps of node degree at five regularly sampled intervals of the developmental trajectories, showing a regionally heterogeneous decrease from young age. B) Definition of local measures of maturation, illustrated on a nonlinearly decreasing trajectory (from the right dorso-lateral pre-frontal cortex). The maximum change in degree Δ*k*_max_ corresponds to the (absolute) difference (decrease or increase) in degree between the maximum and the minimum of the trajectory. The age at minimum degree age(*k*_min_) corresponds to the timing of the minimum of the trajectory. C) Cortical maps of regional maturation measures for trajectories showing evidence of non-zero change (at p_FDR_ < 0.05), predominantly located in association cortex: (i) maximum change in degree, and (ii) age at minimum degree. D) Regions that show greater decreases in degree tend to reach minima of their trajectories later, whether considering all regions (grey) or excluding regions where the trajectory minimum occurs at extrema of the age range (black).

Decreases in node degree were greatest in association cortical areas, such as bilateral dorsolateral prefrontal cortex, medial frontal cortex and supramarginal gyrus, as well as pre-and post-central gyri and several temporal cortical regions. Increases in node degree were less spatially clustered, occurring in isolated nodes within the right cingulate, superior frontal and parietal cortices as well as left cuneus (Fig. 3Ci). Association cortical areas also showed more prolonged decreases in structural correlation, reaching the minimum value of node degree later (Fig. 3Cii). Predictably, it follows that the extent of degree shrinkage Δ*k*_max_ was negatively correlated with the age at which degree reached its minimum value age(*k*_min_), whether considering all regions (Spearman's *ρ* = -0.38, ^p^Spearman < 10^−10, p^perm < 10^−5)^ or excluding regions whose minimum occurred at one of the limits of the age range (Spearman's *ρ* = -0.45, ^p^Spearman < 10^−10^, ^p^perm = < 10^−5;^ Fig. 3D).

Age-related non-linear changes in nodal connection distance (the mean Euclidean distance of all edges connecting a node within the bootstrap-thresholded network) were summarised using analogous parameters to node degree: **Δd_max_**, the difference between maximum and minimum distance; and **age(*D*_min_)**, the age at which nodal connection distance reached its minimum value. Nodes that demonstrated significantly reduced connection distance (p_FDR_ < 0.05) were located in left dorsolateral prefrontal cortex, left supramarginal gyrus and right superior parietal cortex (**SI Fig. S7C**). Decreases in node connection distance were negatively correlated with age at minimum connection distance, whether considering all nodes (Spearman's *ρ* = -0.38, p_Spearman_ < 10^−10^, p_perm_ < 10^−5^) or excluding nodes whose minimum occurs at one of the limits of the age range (Spearman's ρ = -0.25, p_Spearman_ = 0. 0027, p_perm_ = 0.0036) (**SI Fig. S7D**). Finally, decreases in node connection distance were positively correlated with decreases in node degree (Spearman's ρ = 0.32, p_Spearman_ = 1.9 10^−8^, p_perm_ = < 10^−5^) (**SI Fig. 7E**). I n other words, nodes that had the greatest reduction in hubness during adolescence also tended to have the greatest reduction in connection distance.

To contextualise changes in structural network architecture with respect to maturation of cortical morphology, we related regional measures of cortical network development to rates of change of cortical thickness (CT) and magnetization transfer (MT, a measure of myelination), evaluated as the slope of a linear model fitted to the cross-sectional values. The maximum change in node degree was (weakly) positively correlated to the rate of thinning (ΔCT; Spearman's ρ = 0.16, p_Spearman_ = 0.0050, p_perm_ = 0.023; unaffected by excluding three outlier regions which showed ΔCT>0, Spearman's ρ = 0.15, p_Spearman_ = 0.0070, p_perm_ = 0.028; Fig. 4Ai), and more strongly negatively correlated to the rate of intra-cortical myelination (ΔMT; Spearman's ρ = -0.32, p_Spearman_ = 6.6 10^−9^, p_perm_ = 7 10^−4^; Fig. 4Aii). Following Whitaker, Vértes et al. (2016), myelination analyses were conducted at 10 fractional depths between the pial surface and the grey/white matter boundary, as well as two absolute depths into white matter. The strength of association between local adolescent myelination (indexed by ΔMT) and adolescent decrease of node degree (indexed by **Δ*k*_max_**) was greatest when ΔMT was measured at about 70% of cortical depth from the pial surface to the grey/white matter boundary (Fig. 4B).

**Figure 4:**
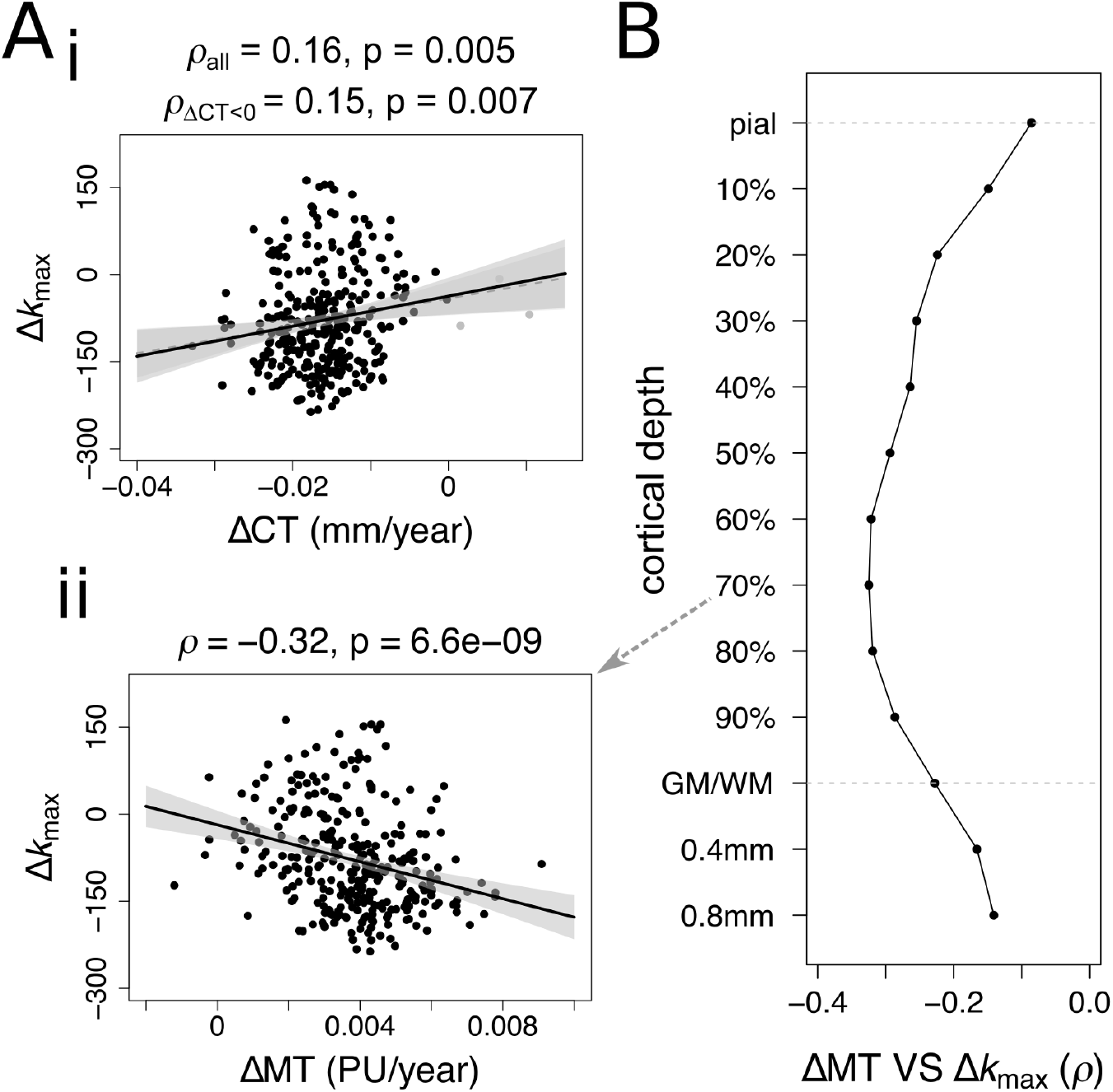
Relationship between maturation of cortical morphology and structural correlation networks. A) Relationship between regional trajectories of cortical morphology and node degree. Maximum changes in nodal degree are only very weakly related to regional rates of i) thinning and ii) myelination (PU = percentage units). The direction of the relationships is such that cortical regions that myelinate more during adolescence are more likely to decrease in node degree and connection distance in the same period. B) Spearman correlation of rate of change myelination to maximal change in degree as a function of cortical depth, including 10 fractional depths from the pial surface to the grey/white matter boundary (GM/WM), as well as two absolute depths into the white matter.

### Age-resolved network changes in relation to the age-invariant network and its communities

Given that most densely connected nodes (hubs) of the age-invariant structural correlation network are predominantly located in association cortex (Whitaker, Vértes et al., 2016), which is also the location of greatest age-resolved decreases in structural correlation, it is not surprising that there is an inverse relationship between age-invariant (weighted) node degree and maximum change in degree (ρ = -0.43, p_Spearman_ < 10^−10^, p_perm_ = < 10^−5^; **SI Fig. S9Bi**). Node degree of the age-invariant network and age at minimum degree were not strongly related (**SI. Fig. S9Bii**).

We further studied adolescent changes in nodal topology in relation to the community structures of the human brain. Many community structures have been proposed to partition the cortex into a set of modules or sub-networks, each comprising a number of functionally and/or anatomically related cortical areas. Here we considered three complementary community structures: (i) the modular decomposition of the age-invariant structural correlation network (7 modules); (ii) the classic von Economo cytoarchitectonic partition of the cortex into classes based on cortical lamination (we used a partition into 7 classes by Vértes et al. (2016), extended from the original partition into 5 classes by von Economo and Koskinas (1925)); and (iii) the prior identification of 7 resting state networks derived from independent components analysis of an independent resting state fMRI dataset (Yeo, Krienen et al., 2011). The three classification systems had similar but not identical community structures; normalised mutual information (NMI, a measure of correspondence between two community structures) ranged from NMI = 0.39 for the relationship between the structural network modules and the resting state fMRI components to NMI = 0.29 for the relationships between both neuroimaging based community structures and the von Economo classification (Fig. 5A).

**Figure 5:**
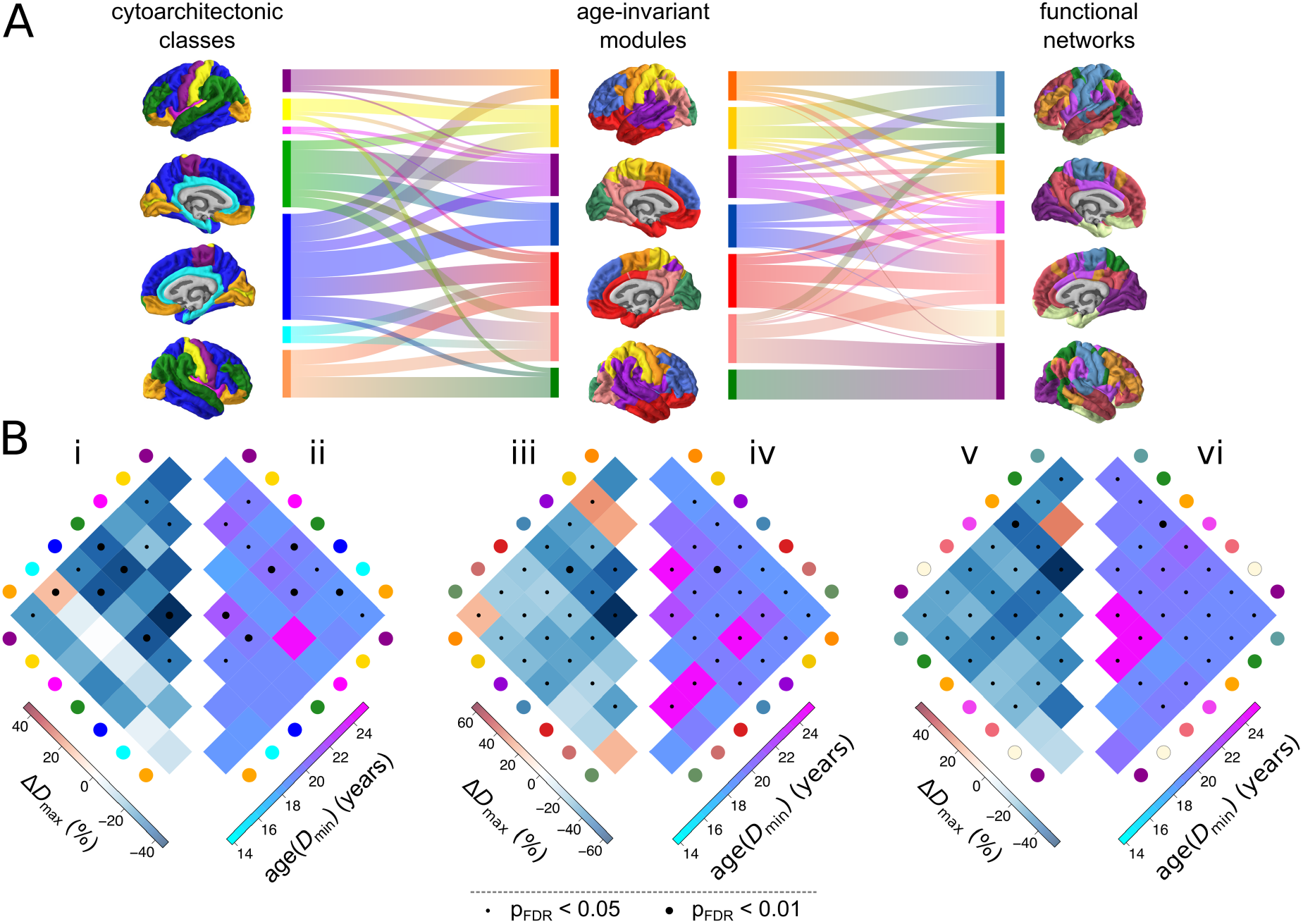
Adolescent development of structural networks in relation to human brain communities. The modular partition used consisted of seven modules, including a parietal "somato-sensory" module (yellow), a frontal "motor" module (orange), an occipital "visual" module (green), an inferior-frontal/temporal module (red), a superior frontal module (blue), a superior temporal/insular module (purple) and a parietooccipital module (pink). A) Comparison of the modular architecture of the age-invariant structural correlation network (middle) to two prior community structures – the von Economo atlas of cytoarchitectonic classes (von Economo & Koskinas, 1925; left) and seven functional intrinsic connectivity networks derived using an independent fMRI data (Yeo & Krienen, 2011; right). The alluvial diagrams between surface plots of community architecture indicate the amount of overlap between individual communities across templates. B) Development of structural correlations within and between corresponding pairs of communities – cytoarchitectonic classes (i-ii), age-invariant modules (iii-iv) and functional intrinsic connectivity networks (v-vi). Left: maximum change in edge density Δ*D*_max_ within and between all pairs of communities. Right: age at minimum edge density age(*D*_min_) within and between all pairs of communities. Dot markers indicate statistical significance of developmental change; small: p_fdr_ < 0.05, large: p_fdr_ < 0.01.

In the context of (i) the age-invariant structural network community structure, the greatest decreases in connection density Δ*D*_max_ were concentrated within the superior frontal module (blue) and within the superior temporal/insular module (purple; Fig. 5Biii); or between the superior frontal module and other modules. The age at minimum density age(*D*_m_¡_n_) tends to occur later within the same modules, as well as the occipito-parietal module (pink; Fig. 5Biv). In the context of (ii) cytoarchitectonic atlas of von Economo & Koskinas (1925), greatest decreases in edge density were concentrated within and between association cortical areas with lamination types 2 and 3 (described as granular isocortex; blue and green respectively) and particularly within class 3 (green; Fig. 5Bi). Association cortical trajectories tended also to reach the age of minimum edge density latest (Fig. 5Bii). In the context of (iii) fMRI resting state networks outlined by Yeo, Krienen et al., (2011), the greatest decreases in edge density were concentrated within the fronto-parietal control network (orange) as well as between this network and the other networks (Fig. 5Bv). Minima of the trajectory are reached latest within the default mode network (salmon red) and the ventral attention network (pink), as well as between these two functional networks (Fig. 5Bvi). In summary, across the three community partitions, the greatest (and latest) decreases in connection density occurred *within* association cortical communities, and (to a lesser extent) *between* those association cortical communities and the remainder of the network.

### Sensitivity analyses

While we had no hypotheses about the shape of the maturational trajectories or the direction of the changes, the finding of a nonlinear decrease in structural correlation (and derived measures of edge density and degree), globally and locally, was somewhat surprising. This is one of the reasons why we conducted numerous sensitivity analyses, to ensure that our findings are not caused or inflated by methodological choices or artefacts.

Our principal findings on bootstrap-thresholded networks were corroborated by similar results from analysis of unthresholded structural correlation matrices (**SI Fig. S6**).

We evaluated robustness of our findings to parameters of the sliding window method, varying the window width and step size over ranges of {40,60,80} and {5,10,20} participants respectively. Results were qualitatively consistent with the above, showing a non-linear decrease in structural correlation both globally and locally (most prominently in association cortex), as well as (weak) relationships of maximum local change in correlation to regional rates of thinning and myelination (**SI Table S2** and **SI Fig. S11**).

Analysis of gender differences failed to identify effects of gender or age-by-gender interactions in the trajectories of structural correlation development (**Supplementary Information**).

We investigated the effect of several potential artefacts, including the presence of regions with low reliability of structural correlations (**SI Fig. S12**) as well as inhomogeneities in the age distribution of participants (**SI Fig. S13**). We found no substantial evidence that the effect of such artefacts could inflate or account for our main finding of a non-linear age-related decrease in structural correlations.

Finally, we investigated whether subtle non-linearities in trajectories of cortical thinning and myelination could be driving non-linearities in trajectories of structural correlation (**SI Fig. S14-16**). Although neither non-linear CT or MT effects are especially strong, subtle nonlinearities in trajectories of cortical myelination appear somewhat more related to structural correlation trajectories than subtle non-linearities in trajectories of cortical thinning.

## Discussion

In the current study we set out to examine the developmental trajectories of human brain structural networks. To this end, we used a novel "sliding window" method of network analysis to resolve age-related changes in human brain structural correlations and probabilistically thresholded brain graphs estimated from MRI data on an age-stratified sample of healthy adolescents and young adults (N=297, aged 14-24 years). We found that global strength of structural correlation and the related topological property of edge density both decreased non-linearly as a function of age: an early phase (14-19.5 years approximately) of rapid decrease in structural correlation was followed by a later phase (20-24 years) of stable or slightly increasing structural correlation. At a regional or nodal level of analysis, cortical areas varied in the magnitude of age-related decrease in nodal degree Δ*k*_max_ and the age at which nodal degree reached its minimum value age(*k*_min_). The 75 cortical areas with significantly decreasing degree tended to mature later, i.e., large negative Δ*k*_max_ was associated with older age(*k*_min_). Further, cortical areas with the greatest shrinkage of degree during adolescence also had the greatest shrinkage of connection distance, i.e., large negative Δ*k*_max_ was associated with large negative Δ*d*_max_. To contextualise these results, we showed that cortical areas with the greatest adolescent changes in brain structural connectivity were anatomically concentrated in regions of association cortex that had fast local rates of increasing intra-cortical myelination; and were topologically concentrated on the edges within frontal communities (von Economo classes 2 and 3 and the functional fronto-parietal control network) and the edges connecting frontal communities to the rest of the network. We propose that these results are consistent with the existence of a developmental window for tuning of association cortical connectivity by a combination of parsimoniously pruning some long distance connections while actively consolidating or myelinating the connections which survive.

### MRI studies of adolescent structural brain network development

Adolescent changes in structural correlation networks have previously been investigated, as pairwise changes across four discrete (non-overlapping) age-bins spanning the range 5-18 years (Zielinski et al., 2010; Khundrakpam et al., 2013). Zielinski et al. (2010) reported largely non-linear changes in the extent of seed-based structural correlation networks. Both the executive control network (seeded in the right dorsolateral prefrontal cortex) and the salience network (seeded in the right frontal insula), showed an increase in spatial extent, quantified as the number of voxels whose grey matter intensity significantly correlated with the seed. Conversely, our approach suggests a decrease in the structural correlation within association areas and related structural, cytoarchitectonic and functional communities. Beyond the difference in methods (voxel-wise seed-based vs. parcel-wise all-to-all regions), this discrepancy could be due to the different morphometric measures used, known to show differences in both trajectories of adolescent maturation (Wierenga et al., 2014; Ducharme et al., 2015), and (age-invariant) structural correlation (Sanabria-Diaz et al., 2010; Yang et al. 2015),. Further, Khundrakpam et al. (2013) reported decreases in regional efficiency of primary sensorimotor regions, alongside increases in regional efficiency of paralimbic and association regions. These results align with our own, through the strong dependence of the properties of graphs thresholded to fixed edge densities (as in Khundrakpam et al. (2013)) on the mean of the correlation distributions from which they were derived. Networks with lower correlations lead to more random topology, exhibiting higher efficiency and lower clustering (Fornito et al., 2013; van den Heuvel et al., 2017). Therefore, our finding of decreases in structural correlation within association cortical areas aligns with reports by Khundrakpam et al. (2013) of increased regional efficiency in these regions. Beyond development of structural networks resolved using distinct age-groups, several studies have investigated coordinated maturation of cortical morphology during adolescence (Raznahan et al., 2011; Alexander-Bloch et al., 2013; Sotiras et al., 2017).

Adolescent development of structural connectivity has also been investigated using diffusion imaging and tractography, although such studies report heterogeneous findings. Lim et al. (2013) showed decreases in structural connectivity from childhood (4 years) to adulthood (40 years), concentrated predominantly on strong tracts, located within modules – which qualitatively agrees with our findings. However, Chen et al. (2013) reported increases in the number of streamlines and edge density from childhood (5 years) to adulthood (30 years). Recently, Baum et al. (2017) reported increases in within-module connectivity, and decreases in between-module connectivity in tractography-derived white matter networks. While tractography-derived structural connectomes show some overlap with structural correlation networks (Gong et al., 2012), interpretation of developmental changes in white-matter connectivity relative to development of structural correlations will require concurrent studies of both modalities in the same datasets. It is worth noting that when grey and white matter structural networks were both constructed using the same method (structural correlation), both showed similar patterns of correlation and similar developmental changes from 7 to 14 years (Moura et al., 2016).

Adolescent development of brain connectivity has also been investigated using fMRI. Early functional connectivity studies have reported increases in the strength of long-range and within-network functional connections (and decreases in the strength of short-range functional connections) (Fair et al., 2009; Supekar et al., 2009; Dosenbach et al., 2010). Later studies have reported qualitatively similar findings, but with attenuated effect sizes following control for the effects of motion (Satterthwaite et al., 2012, 2013). While findings such as increasing within-module functional connectivity may seem to disagree with our findings of decreased within-network structural correlation, these constitute disparate modalities that have not always yielded concomitant results (Fornito & Bullmore, 2015). Beyond studies concurrently investigating adolescent development of structural and functional networks using the same dataset(s), the combination of structural, diffusion and functional MRI data using methods such as multimodal fusion (Calhoun & Sui, 2016), computational modelling (Breakspear, 2017) or morphometric similarity (Seidlitz et al., 2017) might be useful to reconcile findings from diverse modalities.

### Relationship to axo-synaptic connectivity (and its adolescent pruning)

Our results extend previous studies of structural network development (Zielinski et al., 2010; Khundrakpam et al., 2013) by reporting smooth and non-linear trajectories of structural network development during adolescence. The early phase of major decrease in structural correlation, nodal degree, and nodal connection distance could represent loss of anatomical connectivity to association cortical areas. The simplest interpretation is that reduced structural correlation or degree represents pruning of synaptic connections or attenuation of axonal projections. There is a large body of prior evidence in support of the concept of synaptic pruning during adolescence (Huttenlocher & Dabholkar, 1997; Petanjek et al., 2011) and this mechanism has been suggested to explain age-related cortical shrinkage (Tau & Peterson, 2009), which was correlated with age-related degree shrinkage in these data. However, the security of this interpretation rests on the more fundamental assumption that structural correlation measured from MRI data on multiple subjects is a reasonable proxy marker of the average weight of axo-synaptic connectivity between regions (Alexander-Bloch et al., 2013). Beyond humans (Gong et al., 2012), there is evidence of such correspondence from animal models (Yee et al., 2017).

The identification of structural correlation networks in mice (Pagani et al., 2016) suggests that they might encompass general features of cortical architecture. Specifically, up to 35% variance in structural correlation in mice was explained by a combination of tract-tracing-derived structural connectivity, gene expression and distance (Yee et al., 2017), providing a link of the macroscopic structural networks to underlying microscale cortical organisation. The relationship of structural correlation networks to gene expression has also been investigated within humans using the present data, demonstrating overlap between regional co-expression of genes (Hawrylycz et al., 2012), particularly of a subset of genes enriched in supra-granular layers of cerebral cortex, and structural correlation patterns (Romero-Garcia et al., 2017). Moreover, association cortical hubs of the (age-invariant) structural correlation network showed the greatest expression of genes related to synaptic transmission, oligodendroglia as well as schizophrenia, suggesting a potential pathogenic role in abnormal consolidation of association cortical regions (Whitaker, Vértes et al., 2016). Generally, the profound adolescent maturational changes in cortical architecture are thought to underlie the frequent emergence of psychiatric disease in this period, as a result of abnormal development (Paus et al., 2008; Silbereis et al., 2016).

### Adolescent maturation of structural correlation and regional cortical structure

We note that the association of changes in structural network architecture to rates of cortical thinning is relatively weak. Given that (age-invariant) structural correlation networks are thought to emerge as a result of synchronised maturation (thinning) of cortical regions over adolescence (Raznahan et al., 2011; Alexander-Bloch et al., 2013), perhaps the changes in structural correlation might be more closely related to changes in the *rates of change* of cortical thinning, which in a longitudinal dataset were shown to peak in adolescence (Zhou et al., 2015). An additional possible explanation for the adolescent decrease in structural correlation is a "decoherence" related to inter-individual differences in the timing of maturation of association areas – although the verification of such a hypothesis would again require longitudinal data. On a related note, recent work on functional connectivity has shown an adolescent increase the "distinctiveness" of individual functional connectomes (Kaufmann et al., 2017). We further note that the association of changes in structural network architecture to rates of myelination is stronger (than to rates of cortical thinning), and that subtle non-linearities in trajectories of myelination seem more strongly related to nonlinearities in trajectories of structural correlation, suggestive of the idea that myelination may be a driver of (changes) in structural covariance. This could be further investigated through concurrent analysis of (adolescent) changes in structural correlation and white matter architecture.

Generally, the weakness of association between rates of change of morphology (**ΔCT** and **ΔMT**) and structural network architecture (**Δk_max_**) suggests that rates of change of structural network properties explain substantial variation of brain structure with age, above and beyond the rates of thinning and myelination. As an intrinsic regional measure, cortical thickness can be considered less complex than a measure of relationships between regions (across participants) such as structural correlation; however, the biological hierarchy could well be the opposite, whereby cortical thickness and its changes might be a signature of underlying changes in axonal connectivity. This hypothesis could be tested, using invasive studies of concurrent development of axonal connectivity and cortical thickness in model species. In humans, the differential variance contained within cortical morphology and structural network architecture could be investigated through further within-population comparisons of these measures, in (1) their ability to discriminate between case-control populations, (2) their association to behavioural and cognitive measures and (3) their heritability. For example, patients with childhood-onset schizophrenia have shown differences in adolescent trajectories of both cortical thinning (Alexander-Bloch et al., 2014) and structural correlation (Zalesky et al., 2015) relative to healthy controls, but the measures have not been explicitly compared.

Notably, changes in structural network architecture were more strongly related to the rate of myelination (at 70% depth) than the rate of cortical thinning, suggesting that layer-specific intra-cortical myelination might be a more sensitive marker or cortico-cortical connectivity than cortical thickness (assuming, as above, that structural correlation is a marker of connectivity). Moreover, this finding echoes our earlier finding of the rate of myelination being fastest at 70% depth between the pial surface and the grey/white matter boundary, and the relationship between rate of cortical thinning and rate of myelination being strongest at this depth (Whitaker, Vértes et al., 2016). We have previously suggested a link of these changes to histological evidence of greatest rates of myelination at similar cortical depths in rodents (Mengler et al., 2014; Tomassy et al., 2014; Hammelrath et al., 2016).

### Methodological considerations

Recently, a number of studies have pointed out effects of participant motion on the quality of structural MRI scans, including on estimates of regional morphological measures such as cortical thickness (Reuter et al., 2015; Alexander-Bloch et al., 2016; Savalia et al., 2017). While we have carried out stringent quality control of our structural scans and FreeSurfer reconstructions of cortical thickness (details in **Supplementary Information**), we cannot completely rule out potential artefactual effects of motion on our results. Thus, further analysis of structural correlation development in datasets including estimates of head motion from volumetric tracking (Tisdall et al., 2012, 2016) or novel automated estimates of data quality (Shehzad et al., 2015; Pizarro et al., 2016; Rosen, Roalf et al., 2017) will be important in the future.

The estimated changes in structural network organisation are inevitably dependent on parameters of the sliding window method used. The selection of sliding window parameters, including window width and step size (in units of number of participants) involves several trade-offs. On one hand, selecting a wider window increases the robustness of correlations within each of those windows, as they are estimated using more participants; on the other hand, the median ages of participants within each window will cover a narrower portion of the overall age-range. Furthermore, while a smaller step size will provide a greater density of windows and hence time-points for curve fitting and trajectory characterisation, a denser sampling of data will exacerbate issues with the inevitably uneven distribution of subjects across the age-range studied, which in effect corresponds to an unevenly sampled time-series. Future development of tools for the analysis of unevenly sampled time-series (Eckner, 2014) should help alleviate these issues.

Furthermore, depending on the sliding window parameters, relatively few summary data points may be obtained. The subsequent fitting of nonlinear smoothing splines (with up to ~3.5 degrees of freedom) to such scarce data warrants care when interpreting evidence of non-linearity – despite evidence from both the AIC and BIC that smoothing splines provide a better quality of fit than linear models. Still, it is reassuring that trajectories remain consistently nonlinear across bootstrapped samples (within unthresholded correlation networks) and that evidence of a nonlinear trajectory seems more pronounced after bootstrap thresholding. Changes in structural network architecture remain qualitatively consistent in both their spatial location and relationship to changes in morphology when simple, linear models are used. The scarcity of data points may also lead to uncertainties in measures used to characterise the maturational trajectories, including the measures of maximum change and age at minimum of the trajectory. Finally, it remains ambiguous whether the tendency of the global trajectory of structural correlation to slightly increase from the minimum around age 19 towards age 24 years is significant, or whether the trajectory can be seen as levelling-off. It seems reasonable that the few nodes presenting increases in structural correlation (e.g. within right cingulate cortex) would be driving this effect. Thus, until these results are validated in an additional dataset, care is necessary in some aspects of their interpretation.

Further, practical applicability of structural correlation networks is limited by the fact that they represent a group construct. Still, an advantage of structural correlation networks over structural connectomes derived from diffusion imaging using tractography is the relative simplicity of the structural MRI acquisitions compared to diffusion imaging, which in light of its longer acquisition is more prone to motion artefacts (Yendiki et al., 2014), and within which tractography presents considerable challenges (Thomas et al., 2014; Reveley et al., 2015; Maier-Hein et al., 2016). Efforts to derive measures of individual contribution to structural correlation networks (Saggar et al., 2015) or fully individual networks from structural imaging (Tijms et al., 2012; Kong et al., 2014, 2015) including through the combination of multi-modal features (Seidlitz et al., 2017) should increase the practical applicability of structural correlation network research.

In reporting a late maturation of association cortical regions, our results are potentially compatible with the developmental mismatch hypothesis, which proposes that late maturation of prefrontal regions (involved in cognitive control), compared to an earlier development of subcortical regions (implicated in reward processing) results in adolescent increases in risk-taking and sensation-seeking behaviours (Mills, Goddings, Clasen, Giedd, & Blakemore, 2014). However, the verification of such a hypothesis will require the inclusion of both subcortical regions and behavioural data in future analyses.

Finally, structural network architecture is known to mature across the lifespan (DuPre & Spreng, 2017), including during both early childhood (Geng et al., 2016) and late adulthood (Hafkemeijer et al., 2014). Our focused age-range prohibits us from conclusively ascertaining the specificity of these changes to adolescence. For example, extending the analyses presented herein to wider age-ranges would help disambiguate whether the non-linear decreases in structural correlation level off or increase in young adulthood. In general, the wide applicability of the methods used herein should enable investigations of the maturation of structural brain networks, as well as other networks constructed in a similar manner (including for example networks of relationships between psychopathological symptoms; Borsboom & Cramer, 2013), across the life-span.

### Conclusion

During adolescence, human brain structural correlation networks demonstrate a non-linear reduction of connectivity of association cortical areas, predominantly in frontal cortex, that is compatible with a developmental process of pruning combined with consolidation of surviving connections.

## Availability of data and code

Data for this specific paper has been uploaded to the Cambridge Data Repository (https://doi.org/10.17863/CAM.8856) and password protected. Our participants did not give informed consent for their questionnaire measures to be made publicly available, and it is possible that they could be identified from this data set. Access to the data supporting the analyses presented in this paper will be made available to researchers with a reasonable request to. The code used to conduct analyses is available from FV's github: https://github.com/frantisekvasa/structural network development (DOI: 10.5281/zenodo.528674).

## Acknowledgments

We would like to thank Philip T. Reiss and Lisa Ronan for help with the fitting of smoothing splines, Konrad Wagstyl for help with quality control of MRI scans and labelling of regions according to the von Economo atlas, and Manfred Kitzbichler, Håkon Grydeland, Armin Raznahan and Lizanne Schweren for helpful discussions.

## Author contributions

Designed the study: PF, RJD, PBJ, IMG, ETB. Conceived and designed analyses: FV, JS, GR, OS and ETB. Processed and quality controlled data: KJW, RRG, FV and PEV. Conducted analyses: FV. Contributed analysis tools and code: FV, JS, RRG, KJW, PEV, MS and AAB. Wrote the manuscript: FV and ETB. Critically appraised the manuscript: all authors.

